# Basolateral amygdala glutamatergic neurons maintain aversive emotional salience

**DOI:** 10.1101/186072

**Authors:** Auntora Sengupta, Joanna O.Y. Yau, Philip Jean-Richard Dit Bressel, Yu Liu, E. Zayra Millan, John M. Power, Gavan P. McNally

**Author notes:** Correspondence to: Gavan P. McNally PhD School of Psychology UNSW Sydney, NSW Australia.

## Abstract

Basolateral amygdala (BLA) glutamatergic neurons serve a well-accepted role in fear conditioning and fear extinction. However, the specific learning processes related to their activity at different times during learning remain poorly understood. We addressed this using behavioral tasks isolating distinct aspects of fear learning in rats. We show that brief optogenetic inhibition of BLA glutamatergic neurons around moments of aversive reinforcement or non-reinforcement causes reductions in the salience of conditioned stimuli, rendering these stimuli less able to be learned about and less able to control fear or safety behaviours. This salience reduction was stimulus-specific, long-lasting, and specific to aversive emotional states - precisely the goals of therapeutic interventions in human anxiety disorders. Our findings identify a core learning process disrupted by brief BLA optogenetic inhibition. They show that a primary function of BLA glutamatergic neurons is to maintain the salience of conditioned stimuli. This is a necessary precursor for these stimuli to gain and maintain control over fear and safety behavior.

**Significance statement:** The amygdala is essential for learning to fear and learning to reduce fear. However, the specific roles served by activity of different amygdala neurons at different times during learning is poorly understood. We used behavioral tasks isolating distinct aspects of learning in rats to show that brief optogenetic inhibition of BLA glutamatergic neurons around moments of reinforcement or non-reinforcement disrupts maintenance of conditioned stimulus (CS) salience. This causes a stimulus-specific, long-lasting, and aversive emotion specific deficit in the ability of the CS to be learned about or control fear responses. These consequences are the precisely goals of therapeutic interventions in human anxiety disorders. Our findings identify a core learning process disrupted by brief BLA optogenetic inhibition.

---

The capacity to learn about and respond to sources of danger in the environment is essential to survival. Within the laboratory, Pavlovian fear conditioning is most commonly used to model this learning. In conditioning, subjects, typically rats or mice, receive pairings of an initial neutral conditioned stimulus (CS) with an aversive unconditioned stimulus (US) such as footshock. A large body of work shows that the amygdala is essential for Pavlovian fear learning.

The amygdala contributes to both acquisition and expression of Pavlovian fear learning (Davis, 1992; Lüthi and Luscher, 2014; Maren and Quirk, 2004; Pare et al., 2004; Schafe et al., 2001). Within amygdala, glutamatergic neurons, notably CaMKIIα principal neurons of the basolateral amygdala (BLA^CaMKIIα^), are especially important (Sah et al., 2003). BLA^CaMKIIα^ neurons show plasticity during fear conditioning (Marek et al., 2013; Maren and Quirk, 2004; McKernan and Shinnick-Gallagher, 1997). BLA^CaMKIIα^ neurons are subject to complex regulation by multiple families of GABAergic interneurons (Ehrlich et al., 2009; Tovote et al., 2015; Wolff et al., 2014) and are robustly excited by aversive USs, such as footshock (Wolff et al., 2014). This US-evoked activity is essential to causing fear learning. For example, brief BLA^CaMKIIα^ optogenetic inhibition during the shock US impairs fear learning (Namburi et al., 2015; Wolff et al., 2014). Moreover, excitation of BLA^CaMKIIα^ augments fear learning (Johansen et al., 2014) and can serve as a US in its own right, supporting modest responding to a CS that signals such excitation (Gore et al., 2015; Johansen et al., 2010).

However, the specific learning processes related to this US-evoked activity and that is affected by optogenetic manipulation of BLA^CaMKIIα^ neurons during fear conditioning remain unknown. Fear learning involves multiple processes and even the simple delivery or omission of a footshock US has many effects on learning. The primary learning-related functions of the shock US (i.e. reinforcement) or its absence (i.e. non-reinforcement) in fear conditioning are maintenance of CS salience enabling the CS to be learned about and responded to, contribution to a prediction error computation, and updating as well as storing the most recent aversive value of the CS (for review see Hall and Rodríguez, 2017).

BLA^CaMKIIα^ neurons may contribute to many of these functions. However, which functions are linked to BLA^CaMKIIα^ activity during aversive reinforcement and non-reinforcement remains poorly understood.

We addressed this by manipulating BLA^CaMKIIα^ during reinforcement and non-reinforcement in Pavlovian fear conditioning. We examined the impact of BLA^CaMKIIα^ optogenetic inhibition on fear learning, extinction learning, appetitive learning, safety learning, simple discrimination learning, and CS preexposure learning. We show that optogenetic inhibition during moments of aversive reinforcement or non-reinforcement renders a CS less able to be learned about and less able to control fear or safety behaviours. This salience reduction is stimulus-specific, long lasting, and specific to aversive emotional states.

## Materials and Methods

Experimentally naive male Sprague Dawley rats (260-350g) were obtained from Animal Resources Centre (Murdoch, Western Australia. Animals were housed in groups of 4 in plastic cages in ventilated racks in a colony room maintained on a 12:12 light dark cycle (lights on 07:00). Food and water were freely available until 3 days prior to the start of behavioral training when rats were maintained on 65g of standard lab chow (Gordon’s Rat and Mouse food) per day (in addition to any pellets earned) and continuous access to water.

### Apparatus

All behavioural procedures were conducted in eight identical operant chambers with dimensions 24 (length) × 30 (width) × 21cm (height). The top, rear wall and hinged door of the chamber was constructed of Perspex. The sidewalls of the chamber were constructed of stainless steel panels. All chambers had a grid floor constructed of stainless steel rods, 4mm in diameter spaced 15 mm apart. The grid floor was connected to a constant current generator. An external magazine hopper (measuring 5×5cm) was built in to the left side panel and was attached to a pellet delivery system that delivered 45 g grain pellets (Able Scientific, Biotechnology, Western Australia). A lever was mounted 4 cm to the right of the magazine hopper. Each chamber was placed in a larger sound-attenuating box, dimensions 59.5cm (length) × 59 (width) × 48cm (height). A fan was attached to the wall of the sound attenuating box to provide ventilation during behavioural testing and a 28V houselight mounted on the rear wall of the operant chambers. An LED driver with integrated rotary joint (625nm wavelength; Doric Instruments) was suspended above the operant chamber. The auditory CSs were an 85dB 10Hz clicker and 1800Hz, 85dB tone delivered through a speaker attached to rear wall of the chamber. The visual CSs were a 2Hz flashing LED mounted outside the conditioning box on the roof of the sound attenuating chambers and 28V house light turning off. The US was a 0.6mA 0.5 s scrambled footshock delivered to the grid floor All behavioural protocols were controlled through Med-PC software (Med Associates, Vermont).

### Viral Vectors

Adenoassociated viral (AAV) vectors encoding eNpHR3.0 (AAV5-CaMKIIα-eNpHR3.0-WPRE-eYFP (6 × 10^12^vp/ml)) or eYFP (AAV5-CaMKIIα-eYFP (4 × 10^12^vp/ml)) were obtained from UNC Vector Core (University of North Carolina, Chapel Hill, NC) whereas AAVs encoding AAV5-CamKII-GCaMP6f-WPRE-SV40 (4.25 × 10^13^ vp/ml) were obtained from Penn Vector Core (University of Pennsylvania, Philadelphia, PA).

### Stereotaxic surgery

Prior to surgery rats received an intraperitoneal (i.p.) injection of a cocktail of 100mg/ml ketamine (Ketapex, Apex Laboratories, Sydney, Australia) and 0.3ml/kg xylazine (Rompun; Bayer, Sydney, Australia). Once anaesthetised rats were placed in a stereotaxic apparatus (Model 942, Kopf, Tujunga, CA) and shaved to expose the skin surface of the head. Prior to incision, rats received a subcutaneous (s.c.) injection of carprofen (5 mg/kg) and an injection of 0.5% Bupivacaine (Cenvet, Sydney, Australia) just under the surface of incision site. Following incision, metal screws were positioned around the craniotomies and attached to the skull. A hand drill was then used to make two craniotomies above the BLA and a 5μl, 30-gauge conical tipped microinfusion syringe (Hamilton) was used to infuse 0.75 μl of AAV vectors into BLA (A-P -3.00; M-L ± 5.00; D-V -8.60 in mm from bregma) (Paxinos and Watson, 2007) over a 3 min period at a rate of 0.25 μl/minute (UMP3 with SYS4 Microcontroller; World Precision Instruments, Sarasota, FL). The syringe was left in place for 5-7 minutes to permit diffusion of the injected vectors. After viral infusion, fibre optic cannulae were lowered into the BLA (A-P -3.00; M-L ± 5.00; D-V -7.50 for eNpHR3.0, -8.31 mm for gCaMP6f in mm from bregma) (Paxinos and Watson, 2007). These cannulae were held in place by dental cement anchored to the screws and the skull. After surgery, rats were injected i.p. with 0.3 ml of 300mg/ml solution of procaine penicillin (Benicillin; Illium) and subcutaneously with 0.3 ml of a 100mg/ml solution of cephazolin (Hospira, Australia). Daily post-operative and recovery procedures, including weight and infection management, were conducted for the remainder of the experiment. All procedures commenced a minimum of 3 weeks post-surgery.

### Procedures

### Electrophysiology

Rats were deeply anaesthetised with Isoflurane (5%), decapitated and their brain rapidly removed and submerged in ice-cold oxygenated (95% O_2_, 5% CO_2_) sucrose-modified artificial cerebral spinal fluid (ACSF), with the following composition (mM): sucrose, 124; NaCl, 62.5; KCl, 2.5; NaHCO_3_, 26; NaH_2_PO_4_, 1.2; glucose, 10; CaCl_2_, 0.5; and MgCl_2_, 3.3. After 2-3-minutes the brain then mounted to a platform using cyanoacrylate adhesive and submerged in sucrose ACSF. Coronal slices (300μm) were prepared using a vibratome (model VT1200, Leica, Wetzlar, Germany) and then maintained in a holding chamber containing standard ACSF (119 mM NaCl, 2.mM 5 KCl, 1.3 mM MgCl_2_, 2.5 mM CaCl_2_, 1.0 mM Na_2_H_2_PO_4_, 26.2 mM NaHCO_3_, 11 mM glucose, equilibrated with 95% CO2, 5% O2) in a Braincubator (PAYO Scientific, Sydney). Slices were incubated at 32 °C for 30 minutes then maintained at 18 °C until recording.

For electrophysiological recordings, brain slices were transferred to a recording chamber and continuously superfused with ACSF (2 ml min^-1^) at 30° C. Whole-cell patch clamp recordings were made from visually identified neurons using a microscope (Zeiss Axio Examiner D1) equipped with a 20 x water immersion objective (1.0 NA). Patch pipettes (2 – 5 MΩ) were filled with an internal solution containing (in mM) 135 KMeSO_4_, 8 NaCl, 10 HEPES, 2 Mg_2_-ATP, 0.3 Na_3_-GTP, 0.3 EGTA, 0.05 Alexa 594 (pH 7.3 with KOH, 280 – 290 mOsm). Voltage recordings were amplified using a Multiclamp 700B amplifier (Molecular Devices, California, USA), filtered at 6 kHz, digitized at 20 kHz with a Digidata1440A (Molecular Devices) interface, controlled using AxoGraph (Axograph, Sydney, Australia). eNpHR was activated by 535 nm light using an LED fluorescence illumination system (pE-2, CoolLED, Andover, UK). Excitatory postsynaptic potentials (EPSPs) were evoked via a monopolar stimulator fabricated from a patch-pipette filled with ACSF positioned within the BLA. Stimuli were supplied by a constant voltage isolated stimulator (DS2A, Digitimer Ltd, Hertfordshire, England). Only cells that had resting potentials more negative than -55 mV, action potential amplitudes > 100 mV, and input resistances > 60 MΩ, were considered healthy and included in the data set. Projection neurons were distinguished from local circuit interneurons based on their action potential half-width (> 0.7 ms), their relatively small fast AHP (< 15 mV), and their frequency-dependent spike broadening. Data were not corrected for liquid junction potentials.

### Baseline Lever Pressing

We used conditioned suppression of lever pressing as our measure of learned fear (Sengupta et al., 2016; Yau and McNally, 2015). Rats were trained to lever press to establish a stable baseline lever pressing response. On Days 1 and 2 rats received magazine training in which every lever press was rewarded with the delivery of a pellet. In addition, rats also received free pellet deliveries on a fixed interval (FI) 300 s schedule, in which a pellet was delivered on average every 300 s. Magazine training sessions were terminated after 60 min, or if the rat reached 100 lever presses. On Day 3, rats underwent a 60 min session of lever press training under a variable interval (VI) 30 s schedule. From Day 4 until the end of the experiment, rats were maintained on a VI120 s. These sessions lasted 120 min unless otherwise noted. On days 9-10 rats received pre-exposure to the CSs to be used in the experiment during a 120 min pre-exposure session. Rats were habituated to the tethering procedure for 4 days on Days 7 - 10.

### Fibre Photometry

Rats underwent fear conditioning on Days 11-13 where they received differential fear conditioning to two CSs. One CS (CS+: a 60 s flashing (2Hz) LED light or a 60 s 80-dB clicker, counterbalanced) was paired four times with a 0.6 mA, 0.5 s footshock whereas the other CS (CS-: a 60 s flashing (2Hz) LED light or a 60 s 80-dB clicker, counterbalanced) was not. The trials were presented in a pseudorandom order with an ITI of 1200 to 1800s.

Recordings were completed using Fibre Photometry Systems from Doric and Tucker-Davis Technologies (RZ5P, Synapse). Excitation light was emitted from Doric 465 nm and 405 nm LEDs, controlled via dual channel programmable LED drivers. These wavelengths were reflected into the 400um pre-bleached patch cable via a Doric Dual Fluorescence Mini Cube. Light intensity at the tip of the patch was set to 200-400uW and was kept constant across sessions. At the start of each recording session, patch cables were attached to the fibre optic implant and recording was initiated. Excitation light was delivered 3 minutes before each CS onset and terminated 1 minute after each CS offset.

GCaMP6f fluorescence was collected from the same implant/patch cable. Ca^2+^ dependent (525nm) and isobestic control (430nm) fluorescence signals (corresponding to 465nm and 405nm excitation, respectively) were relayed to femtowatt photoreceivers (Newport, 2151) through dichroic mirrors and bandpass filters within the Doric Fluorescence Mini Cube. Synapse software controlled and modulated excitation light (465nm: 220Hz, 405nm: 330Hz) and demodulated transduced fluorescence signals in real-time (1kHz sampling rate) via the RZ5P. Synapse/RZ5P also received Med-PC signals to record behavioural events and experimenter-controlled stimuli in real time.

### BLA^CaMKIIα^ neurons and fear learning

Rats were attached to bilateral patch cables outputting at least 9mW of 625 nm light, measured through an unimplanted fibre optic cannula, prior to session commencement in this and subsequent experiments. On Days 11-13, rats underwent 120 min conditioning sessions where they received four presentations of 60 s auditory CS (85dB Clicker) coterminating with a 0.5 s 0.6mA footshock US. 625 nm light delivery coincided with US presentations and extended for a further 4.5 seconds post US. There was a randomised inter-trial interval (ITI) of 1200 – 1800 s. For the offset control, rats were trained identically to the above with the single exception that 625 nm light delivery occurred randomly in the ITI between CS-US pairings during conditioning. All rats were tested in a single 70 min session on Day 14. Rats received four 60 s presentations of the auditory CS with an ITI of 900 s.

### BLA^CaMKIIα^ neurons and fear extinction learning

Rats underwent fear conditioning on Days 11-13 where they received four presentations of an auditory CS (85dB Clicker) co-terminating with a 0.5s, 0.6mA footshock US. All CSs were presented with a randomised ITI of 1200 – 1800 s. Fear acquisition sessions were identical to the above. On Days 14-16 rats were extinguished. All rats received four 60 s presentations of the auditory CS which co-terminated with 0.5 seconds of photoinhibition. This extended for a further 4.5 seconds beyond the CS. These stimulus events were presented with a randomised ITI of 490-1200 s. For the offset control, the same offset control rats from the acquisition experiment were trained to fear a different CS (visual CS 4 pairings a day for 3 days with a .5s, 0.6mA footshock US). They were extinguished to the CS in a manner identical to the above with the single exception that 625 nm light delivery occurred randomly in the ITI between CS presentations.

#### Reacquisition of fear conditioning

On Days 17-20 rats that underwent fear reacquisition received four presentations of the auditory CS co-terminating with a 0.5 s, 0.6mA footshock US in 70 min reacquisition sessions. The ITI was randomised 1200 - 1800 s. There was no photoinhibition.

#### Acquisition of appetitive conditioning

On days 17-20 rats that underwent the appetitive conditioning procedure received four presentations of the auditory CS (30 s in duration) followed by the delivery of 3 × 45 mg sucrose pellets in a 60 min session. The lever which had been used for conditioned suppression was retracted during these sessions. The ITI was randomised from 300 - 600s. There was no photoinhibition.

#### BLA^CaMKIIα^ neurons and safety learning

For this experiment baseline lever press training was conducted as described earlier with one exception. The house light was locked on prior to session commencement and remained on after termination of the session because house light off was used as CS in this experiment. On days 9-10 all rats underwent pre-exposure training where they received three 60 s presentations of each of the following stimuli: a flashing LED, the house light turning off, a 85dB clicker, and a 1800Hz, 85dB tone. Each stimulus was presented with a randomized ITI ranging from 600 - 900 s. On days 11-15, rats received discrimination training where the aforementioned visual cues were counterbalanced as CS A and B, while the auditory cues were counterbalanced as CS X and Y. These rats received four 60 s presentations of AX coterminating with a 0.5 s, 0.6mA shock (AX+) and four non-reinforced presentations of BX-per day. These stimuli were presented with randomized ITIs of 600 – 900 s. Photoinhibition was delivered on BX‐ trials only, coterminating for 0.5 s of the BX‐ trial and extending for a further 4.5 seconds. On days 16-17, rats received four 60 s presentations of Y coterminating with a 0.5 s, 0.6mA shock. These stimulus events were presented with a randomized ITI of 1200 – 1800 s. All rats then underwent tests of summation and retardation. On day 18 rats underwent a summation test where they received four 60 s presentations of the compound AY and BY, each. On days 19-21, rats underwent a retardation test where they were conditioned to B via four pairings with a 0.5 s, 0.6mA shock.

An eYFP - naive group was used as a control during the retardation. These rats received the same lever press training but then received simple A+/B-discrimination training. All other behavioral procedures remained the same.

### BLA^CaMKIIα^ neurons and CS preexposure

Rats underwent fear conditioning training on Days 11-14 where they received two presentations of visual CSA (flashing LED) co-terminating with a 0.5 s 0.6mA footshock US and four presentations of an auditory CSX (85dB Clicker) co-terminating with photoinhibition at 0.5 seconds and extending for a further 4.5 seconds. All CSs were presented with a randomised ITI of 600 - 900 s. These training sessions lasted 120 min. On Days 15-16 all rats received two 60 s presentations of a compound CSAX and two presentations of CSA per session, in a counterbalanced order, with a randomised ITI of 490-1200 s. This session lasted 70 min. On Days 17-19 all rats received four presentations of CSX per session with a randomised ITI of 1200 – 1800 s. Each presentation of CSX co-terminated with a 0.5 second 0.6mA foot-shock US. This session lasted 120 minutes.

### Histology

Cannula and viral placements were verified at the end of behavioural procedures. Following transcardial perfusion with 0.9% Saline, 1% sodium nitrite solution and 360μl/L heparin, then 4% buffered paraformaldehyde (PFA), brains were post fixed then placed in 20% hypertonic sucrose solution for 24-48hrs. A cryostat (Leica Microsystems) was used to collect 40μm coronal sections, preserved in phosphate buffered azide (0.1% sodium azide) at 4°C prior to immunohistochemistry. eYFP labelling using single color peroxidase immunohistochemistry.

Free floating sections were washed for 30 min in 0.1M phosphate buffer (PB; pH 7.4), followed by 50% ethanol (EtOH) for 30 min and then 3% hydrogen peroxide diluted in 50% EtOH for 30 min. Sections were blocked (30 min with 5% NHS diluted in PB), then placed in 1:2000 chicken anti-GFP (Life Technologies; A10262) diluted in 0.3% Triton-X, 2% NHS and 0.1M PB (pH 7.4), and incubated at 4°C for 48hrs. Sections were washed three times for 20 min each (PB, pH 7.4), incubated in 1:3000 biotinylated donkey anti-chicken (Jackson ImmunoResearch Laboratories; 703 065 155) (diluted in a solution of 0.3% Triton-X, 2% NHS and 0.1M PB (pH 7.4)), overnight at room temperature. They were then washed in PB (pH 7.4) and incubated in Avidin-Biotin (ABC reagent; Vector Elite kit 6μl/ml avidin and 6μl/ml biotin) diluted with PB containing 0.2% triton (pH 7.4). eYFP-IR was identified using a 3,3 diaminobenzidine tetrahydrochloride hydrate (DAB; D5637-56, Sigma) reaction. Immediately prior to this, sections were washed twice in PB (pH 7.4) and once in 0.1M acetate buffer (pH 6.0). Sections were incubated in DAB solution (0.025% DAB, 0.04% ammonium chloride and 0.2% d-glucose in 0.1M acetate buffer (pH 6.0)). The peroxidase reaction was catalysed by 0.2μl/ml of glucose oxidase and then were rinsed with 0.1M acetate buffer. Fibre placements and extent of eYFP-IR per rat was verified and photographed using a transmitted light microscope (Olympus BX51). Only animals with fibre placements and AAV expression in basolateral amygdala were included in the analyses. 28 rats in total were excluded due to these criteria (1 from photometry, 9 from fear acquisition experiment, 6 from extinction experiment, 6 from safety learning experiment, 6 from simple discrimination experiment).

### Quantification and Statistical Analyses

Suppression ratios were calculated as *SR = a/(a+b)* (Annau and Kamin, 1961), where *a* represents the number of lever presses during the CS period and *b* represents the number of lever presses recorded 60 seconds pre CS. A SR of 0.5 indicates low suppression or low fear while an SR of 0 indicates complete suppression or high fear. For the appetitive conditioning procedure, a ratio of *a/(a+b)*, where *a* represents the number of magazine entries during the CS period and *b* represents the number of magazine entries 30 s prior to the CS, was used. In this instance, a ratio of 1 indicates exclusive entry into the magazine port during the CS, while a ratio of 0.5 indicates equivalent port entries during the CS and the preCS. These data were analysed via orthogonal contrasts (Harris, 2004). The Hay’s Decision-wise error rate (α) was controlled at 0.05 for each contrast.

For fibre photometry, event-related GCaMP6f fluorescence signals were analysed using customized MATLAB scripts. Demodulated fluorescence signals were Gaussian filtered (to smooth each signal) and converted into a movement-corrected ΔF/F signal by applying a least-squares linear fit to the 405nm signal, aligning it to the 465nm signal. This fitted-405 signal was used to determine the Ca2+ dependent signal as follows: ΔF/F = (465 signal – fitted-405 signal)/fitted-405 signal. For each session, ΔF/F signals around behavioural events (time 0) were isolated and aggregated; 4 to 2 s before each event was used as a baseline (set to 0%) to normalise peri-event ΔF/F, the 2 s leading up to each event was defined as the pre-event transient, and the 5 s following each event was defined as the event transient. Area under the curve (AUC) for event transients were calculated by approximating the integral (trapezoidal method) of the isolated normalised ΔF/F curves. AUCs were analysed using single means t-tests against 0.

## Results

### BLA^CaMKMα^ photoinhibition impairs fear learning

Conditioned suppression of lever pressing for food was used to assess conditioned fear. This is a well-established measure of learned fear. It has several advantages. Suppression has a non-zero baseline because rats lever press for a pellet reward at a constant rate, so revealing decreases and increases in fear; there are high levels of activity during training and testing; it is equally sensitive to visual and auditory CSs despite these CSs eliciting different amounts of freezing on the same trial (Bevins and Ayres, 1991); and, its assessment is completely automated.

We used fibre photometry to examine shock US-evoked activity of BLA^CaMKIIα^ neurons. Rats received differential fear conditioning so that one CS (CS+ [visual or auditory]) was paired with a footshock US and a second CS (CS‐ [auditory or visual]) was not. AAV vectors were used to express the genetically encoded calcium (Ca^2+^) indicator gCaMP6f in BLA^CaMKIIα^ neurons (**Figure 1a**) and fibre photometry (n = 6) (Gunaydin et al., 2014) was used to record Ca^2+^ transients during the shock US.

**Figure 1.**
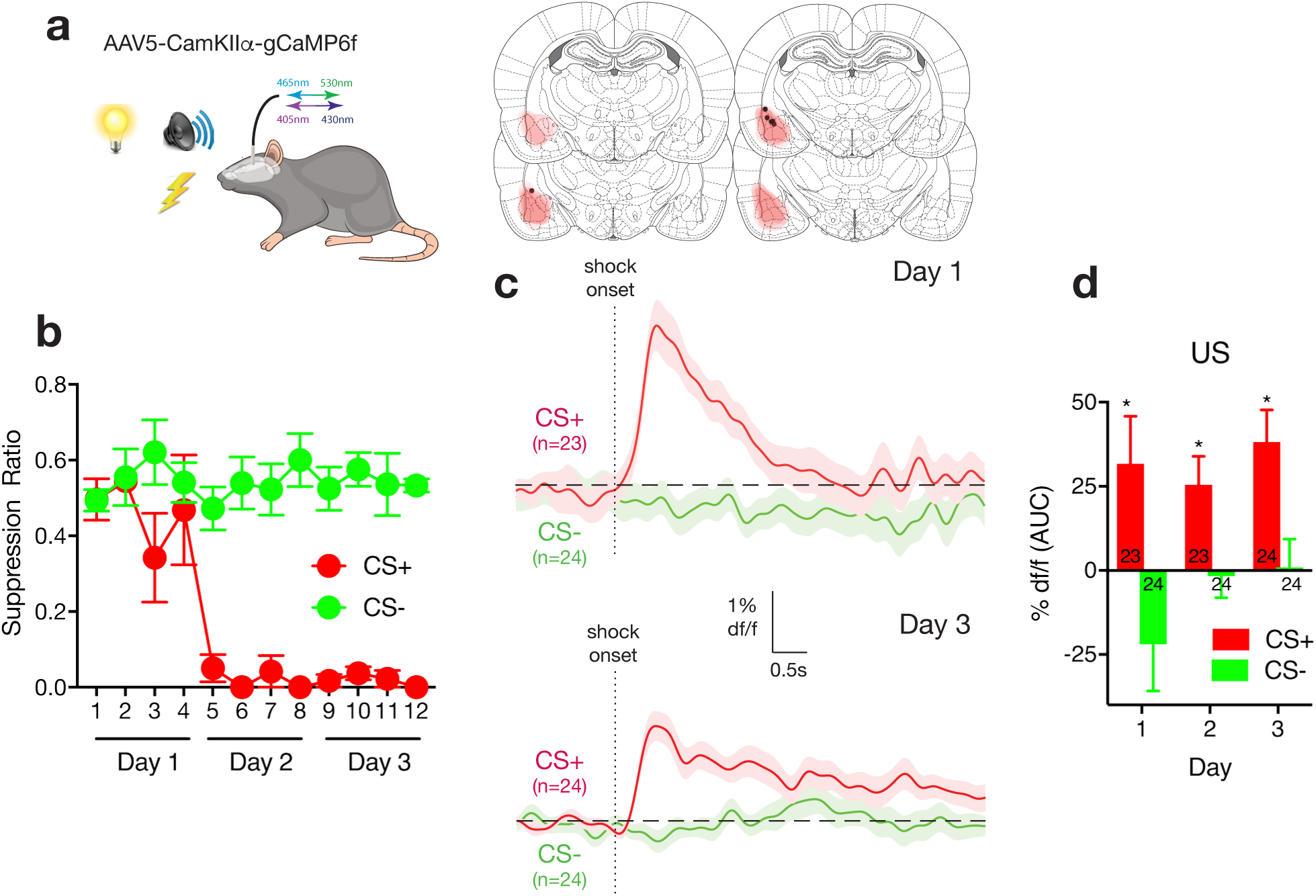
Fibre photometry of BLA^CaMKIIα^ neurons during fear learning. **A,** AAV-CamKIIα-gCaMP6f was applied to BLA. AAV expression across all animals with each rat represented at 10% opacity. **B**, Rats received differential fear conditioning (CS – shock [CS+] CS – no shock [CS-]) and learned to fear CS+ but not CS-. **C**, Ca^2+^ transients during shock US delivery on CS+ trials and omission on CS‐ trials on days 1 and 3 of conditioning. *n* refers to number of trials. **D**, Areas under the ΔF/F curve for the 5 s from onset of shock or shock omission across conditioning. *n* refers to number of trials.

Rats discriminated the dangerous CS+ from the CS-. A suppression ratio of 0.5 indicates no fear and a suppression ratio of 0 indicates high fear. There was selective suppression of lever pressing, indicative of learned fear to the CS+ (**Figure 1b**) (main effect CS+ v CS‐ *F*_(1,5)_ = 86.84, p < .0001; main effect trial *F*_(1,5)_ = 25.33, p = .004; CS x trial interaction: *F*_(1,5)_ = 58.97, p = .001). The shock US evoked robust Ca^2+^ transients on CS+ trials that were absent on CS‐ trials (**Figure 1c**). Area under the curve analyses for the 5 s following the shock US (**Figure 1d**) showed increases for CS+ (Day 1: *t*_22_ = 2.26, p = .03; Day 2: *t*_22_ = 3.01, p = 0.006; Day 3: *t*_23_ = 4.01, p = .0005) but no changes for CS‐ (Day 1: *t*_23_ = -1.58, p = .12; Day 2: *t*_23_ = -0.26, p = 0.79; Day 3: *t*_23_ = .06, p = .95), confirming robust US-evoked activity in BLA^CaMKIIα^ neurons during our fear conditioning preparation.

Next, in separate animals, whole-cell patch-clamp recordings were made from eNpHR3.0 expressing BLA^CaMKIIα^ neurons *in vitro* to confirm photoinhibition. These neurons had the properties of BLA^CaMKIIα^ projection neurons (**Table 1; Figure 2a**). Photoinhibition induced a rapid onset hyperpolarisation that persisted for duration of the light (**Figure 2b**; n = 7 of 7 cells). This was sufficient to inhibit action potentials evoked by depolarising current injections (**Figure 2c**) or synaptic stimulation (**Figure 2d**). Light-evoked responses were not observed in non-eNpHR3.0 expressing neurons.

**Table 1.** Properties of BLA^CaMKIIα^ neurons during whole-cell patch-clamp recordings.

| Property | Mean ± SD | n |
| --- | --- | --- |
| Resting membrane potential | $-69 \pm 7$ mV | 7 |
| Input resistance | $145 \pm 48$ MΩ | 7 |
| Membrane time constants | $29 \pm 13$ ms | 7 |

**Figure 2.**
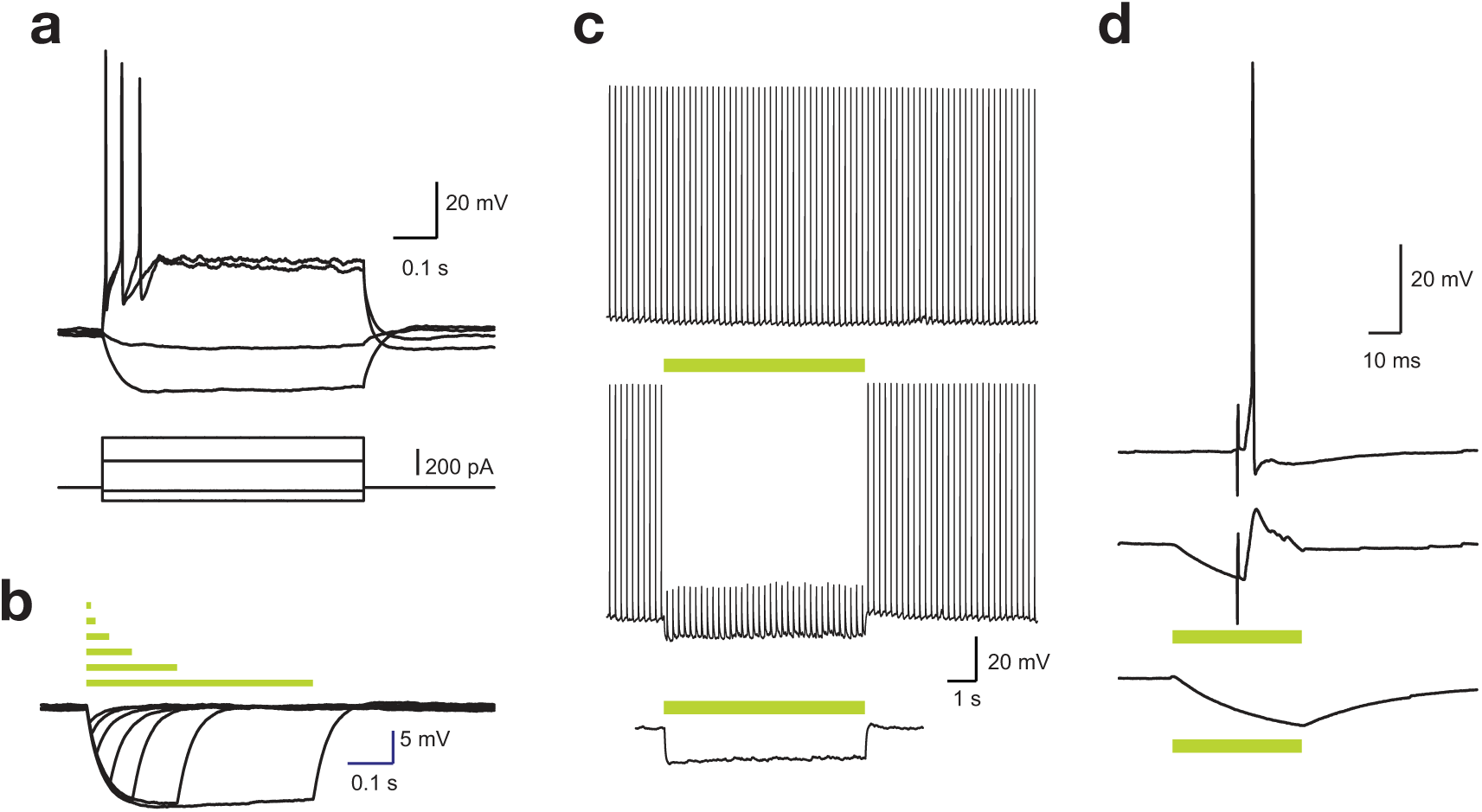
Characterization of BLA^CaMKIIα^ neurons. **A**, Typical voltage response of eNpHR3.0 positive neurons to positive and negative current injections. **B**, Voltage responses to photoinhibition of various durations. Bars indicate timing of the stimuli. **C**, Action potential train evoked by brief current injections (5 ms, 10 Hz) plotted above the response to a 7s light pulse delivered during the AP train, and the response to a 7s light pulse in the absence of depolarizing current injections. **D**, Response to synaptic stimulation (upper), synaptic stimulation during photoinhibition (middle), and photoinhibition alone (lower).

To examine impact of photoinhibition during aversive reinforcement, rats expressing AAV-CaMKIIα-eNpHR3.0-eYFP (CaMKIIα-eNpHR3.0) (n = 7) or AAV-CaMKIIα-eYFP (CaMKIIα-eYFP) (n = 7) in BLA (**Figure 3a**) received fear conditioning to an auditory CS. They received photoinhibition during US delivery and for 4.5 s seconds afterwards (**Figure 3b**), to encompass any activity changes persisting beyond the US. There were no differences between groups in pre-CS lever pressing rates (**Table 2**). Group eNpHR3.0 learned less fear than group eYFP (**Figure 3b**) (main effect of group *F*_(1,12)_ = 4.74, *p* = .05; main effect of day - *F*_(1,12)_ = 294.42, *p* < .0001; no group × day interaction *F*_(1,12)_ = 4.08, *p* > .05). They also expressed less fear on test (*F*_(1,12)_ = 7.89, *p* = .016) (**Figure 3b**), showing that photoinhibition impaired Pavlovian fear learning in conditioned suppression.

**Table 2.**
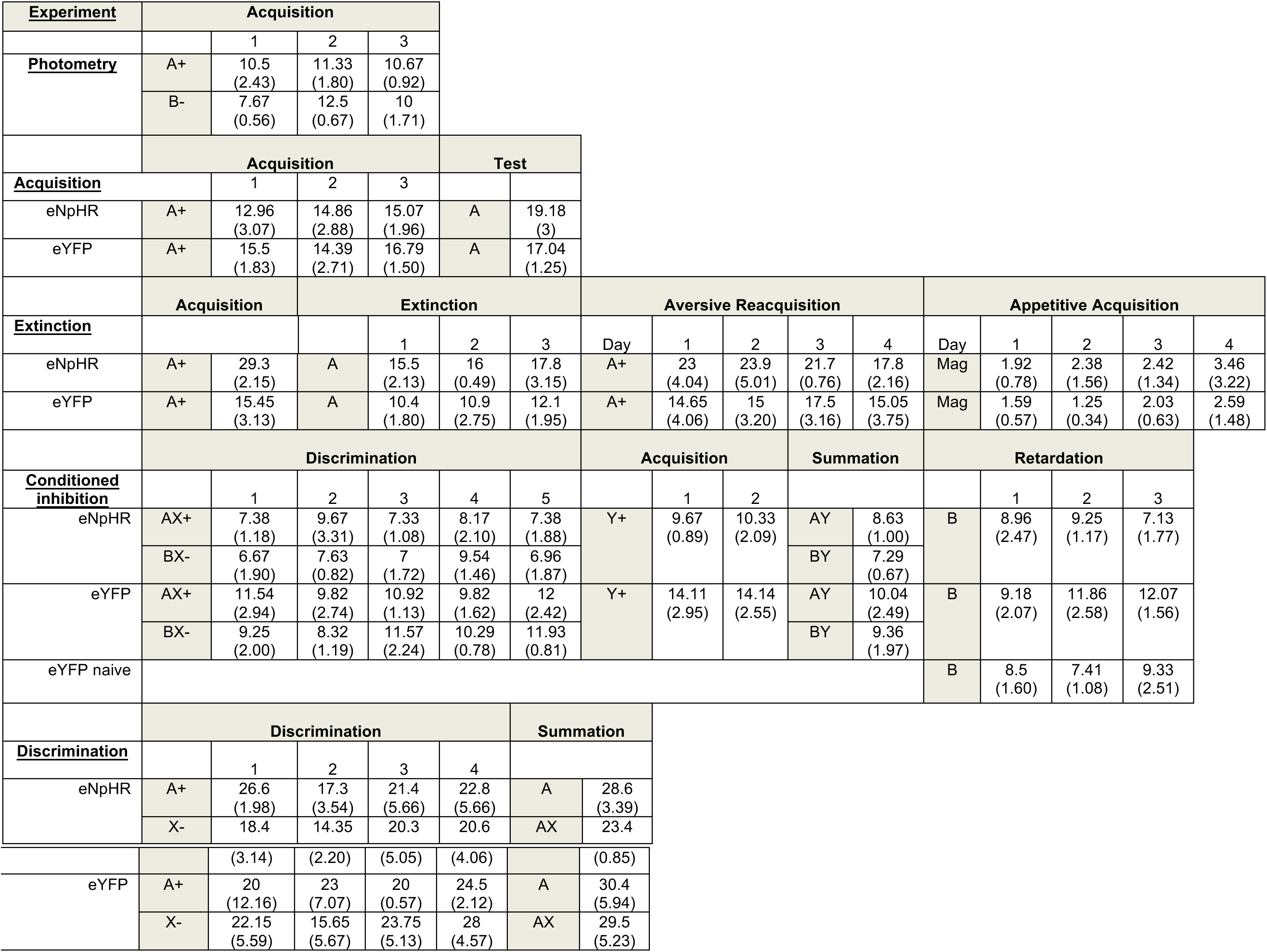
Mean ± SEM pre-CS lever press rates for each group in each experiment.

| Experiment | Acquisition |  |  |  |  |  |  |  |  |  |  |  |  |  |  |  |
| --- | --- | --- | --- | --- | --- | --- | --- | --- | --- | --- | --- | --- | --- | --- | --- | --- |
|  |  | 1 | 2 | 3 |  |  |  |  |  |  |  |  |  |  |  |  |
| Photometry | A+ | 10.5<br>(2.43) | 11.33<br>(1.80) | 10.67<br>(0.92) |  |  |  |  |  |  |  |  |  |  |  |  |
|  | B- | 7.67<br>(0.56) | 12.5<br>(0.67) | 10<br>(1.71) |  |  |  |  |  |  |  |  |  |  |  |  |
|  | Acquisition |  |  |  |  | Test |  |  |  |  |  |  |  |  |  |  |
| Acquisition |  | 1 | 2 | 3 |  |  |  |  |  |  |  |  |  |  |  |  |
| eNpHR | A+ | 12.96<br>(3.07) | 14.86<br>(2.88) | 15.07<br>(1.96) | A | 19.18<br>(3) |  |  |  |  |  |  |  |  |  |  |
| eYFP | A+ | 15.5<br>(1.83) | 14.39<br>(2.71) | 16.79<br>(1.50) | A | 17.04<br>(1.25) |  |  |  |  |  |  |  |  |  |  |
|  | Acquisition |  | Extinction |  |  |  | Aversive Reacquisition |  |  |  |  | Appetitive Acquisition |  |  |  |  |
| Extinction |  |  |  | 1 | 2 | 3 | Day | 1 | 2 | 3 | 4 | Day | 1 | 2 | 3 | 4 |
| eNpHR | A+ | 29.3<br>(2.15) | A | 15.5<br>(2.13) | 16<br>(0.49) | 17.8<br>(3.15) | A+ | 23<br>(4.04) | 23.9<br>(5.01) | 21.7<br>(0.76) | 17.8<br>(2.16) | Mag | 1.92<br>(0.78) | 2.38<br>(1.56) | 2.42<br>(1.34) | 3.46<br>(3.22) |
| eYFP | A+ | 15.45<br>(3.13) | A | 10.4<br>(1.80) | 10.9<br>(2.75) | 12.1<br>(1.95) | A+ | 14.65<br>(4.06) | 15<br>(3.20) | 17.5<br>(3.16) | 15.05<br>(3.75) | Mag | 1.59<br>(0.57) | 1.25<br>(0.34) | 2.03<br>(0.63) | 2.59<br>(1.48) |
|  | Discrimination |  |  |  |  |  | Acquisition |  |  | Summation |  | Retardation |  |  |  |  |
| Conditioned inhibition |  | 1 | 2 | 3 | 4 | 5 |  | 1 | 2 |  |  |  | 1 | 2 | 3 |  |
| eNpHR | AX+ | 7.38<br>(1.18) | 9.67<br>(3.31) | 7.33<br>(1.08) | 8.17<br>(2.10) | 7.38<br>(1.88) | Y+ | 9.67<br>(0.89) | 10.33<br>(2.09) | AY | 8.63<br>(1.00) | B | 8.96<br>(2.47) | 9.25<br>(1.17) | 7.13<br>(1.77) |  |
|  | BX- | 6.67<br>(1.90) | 7.63<br>(0.82) | 7<br>(1.72) | 9.54<br>(1.46) | 6.96<br>(1.87) |  |  |  | BY | 7.29<br>(0.67) |  |  |  |  |  |
| eYFP | AX+ | 11.54<br>(2.94) | 9.82<br>(2.74) | 10.92<br>(1.13) | 9.82<br>(1.62) | 12<br>(2.42) | Y+ | 14.11<br>(2.95) | 14.14<br>(2.55) | AY | 10.04<br>(2.49) | B | 9.18<br>(2.07) | 11.86<br>(2.58) | 12.07<br>(1.56) |  |
|  | BX- | 9.25<br>(2.00) | 8.32<br>(1.19) | 11.57<br>(2.24) | 10.29<br>(0.78) | 11.93<br>(0.81) |  |  |  | BY | 9.36<br>(1.97) |  |  |  |  |  |
| eYFP naive |  |  |  |  |  |  |  |  |  |  |  | B | 8.5<br>(1.60) | 7.41<br>(1.08) | 9.33<br>(2.51) |  |
|  | Discrimination |  |  |  |  | Summation |  |  |  |  |  |  |  |  |  |  |
| Discrimination |  | 1 | 2 | 3 | 4 |  |  |  |  |  |  |  |  |  |  |  |
| eNpHR | A+ | 26.6<br>(1.98) | 17.3<br>(3.54) | 21.4<br>(5.66) | 22.8<br>(5.66) | A | 28.6<br>(3.39) |  |  |  |  |  |  |  |  |  |
|  | X- | 18.4 | 14.35 | 20.3 | 20.6 | AX | 23.4 |  |  |  |  |  |  |  |  |  |

|  |  |  |  |  |  |  |  |
| --- | --- | --- | --- | --- | --- | --- | --- |
|  |  | (3.14) | (2.20) | (5.05) | (4.06) |  | (0.85) |
| eYFP | A+ | 20<br>(12.16) | 23<br>(7.07) | 20<br>(0.57) | 24.5<br>(2.12) | A | 30.4<br>(5.94) |
|  | X- | 22.15<br>(5.59) | 15.65<br>(5.67) | 23.75<br>(5.13) | 28<br>(4.57) | AX | 29.5<br>(5.23) |

**Figure 3.**
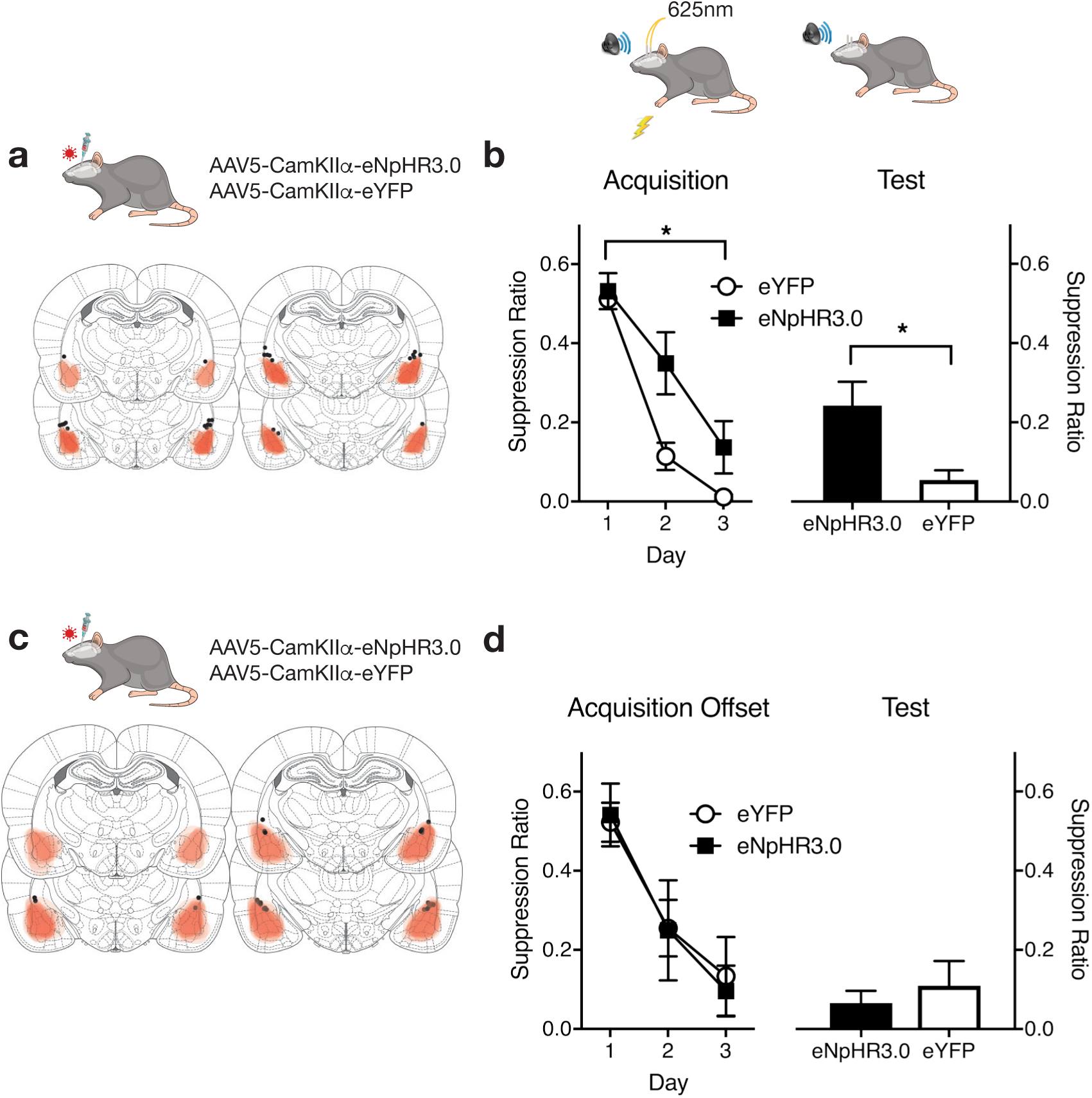
BLA^CaMKIIα^ neurons and fear learning. **A**, AAV expression across all animals with each rat represented at 10% opacity. **B**, eYFP and eNpHR3.0 groups received CS - shock pairings with photoinhibition during the shock US. Photoinhibition impaired fear conditioning. A suppression ratio of 0.5 indicates no fear and a suppression ratio of 0 indicates high fear. **C**, AAV expression across offset control animals with each rat represented at 10% opacity. **D**, Control eYFP and eNpHr3.0 groups received CS – shock pairings and photoinhibition randomly during the inter-trial interval. All data are mean ± SEM and ns are listed in the main text. * p < .05

This impairment was specific to inhibition of US-evoked activity. Separate groups expressing CaMKIIα-eYFP (*n*=7) or CaMKIIα-eNpHR3.0 (*n*=5) in BLA (**Figure 3c**) received fear conditioning with photoinhibition at random times during the intertrial interval. There was no effect on fear learning (acquisition main effect of day *F*_(1,10)_ = 90.48, *p* < .0001, no main effect of group *F*_(1,10)_ < 1, *p* > .05 or group × day interaction *F*_(1,10)_ < 1, *p* > .05) (test: *F*_(1,10)_ < 1, *p* > .05) (**Figure 3d**).

### BLA^CaMKIIα^ photoinhibition augments fear loss during extinction

Just as rats learn to fear a CS that signals shock, so too do they learn to reduce fear to that CS when it is presented in the absence of shock during extinction training. We asked whether BLA^CaMKIIα^ photoinhibition during non-reinforcement would affect fear extinction learning. Rats expressing CaMKIIα-eYFP (*n*=14) or CaMKIIα-eNpHR3.0 (*n*=11) in BLA (**Figure 4a-c**), received fear conditioning to an auditory CS followed by extinction training involving CS alone presentations and photoinhibition at the time of the expected but absent footshock US.

**Figure 4.**
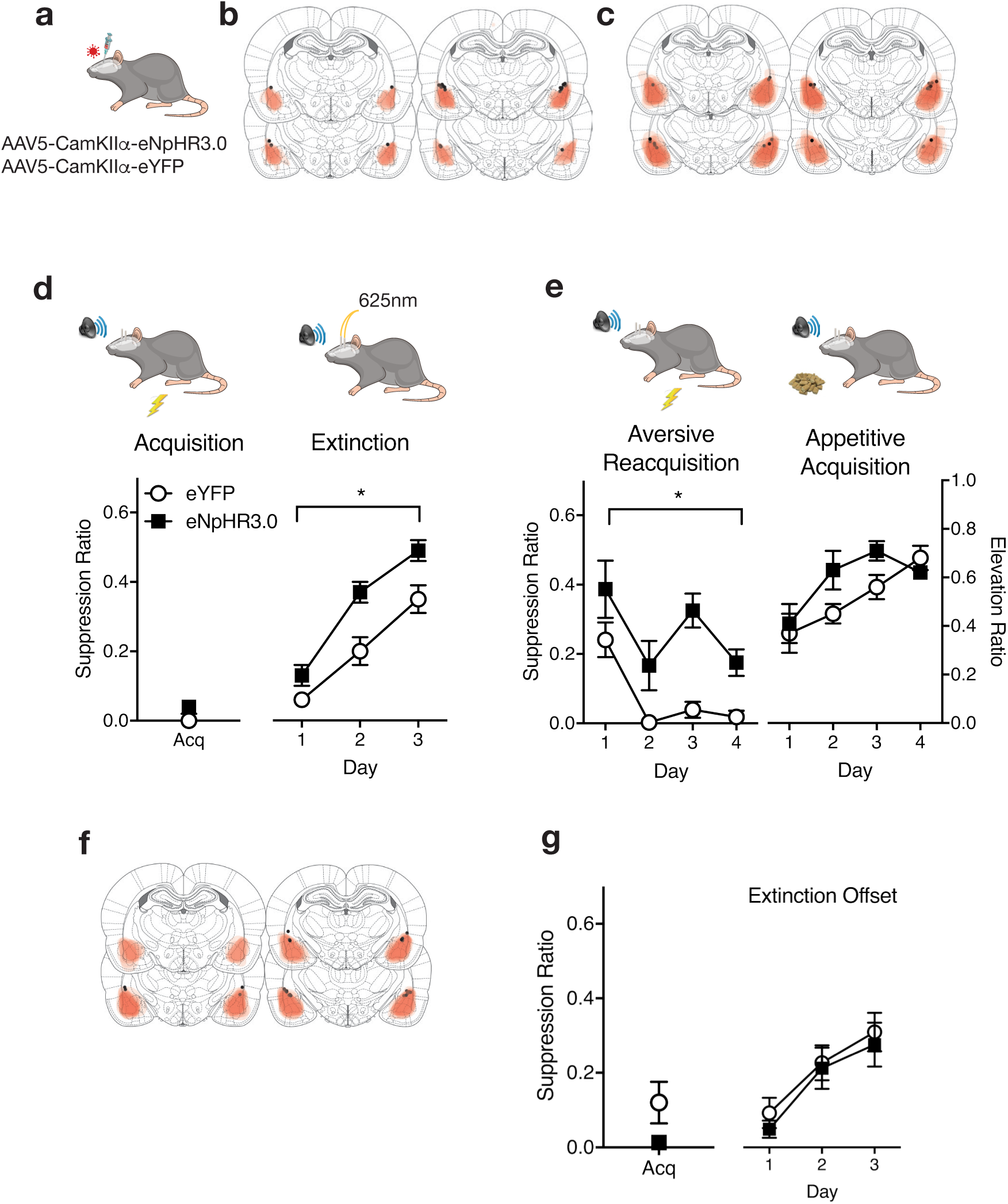
BLA^CaMKIIα^ neurons and fear extinction learning. **A**, AAV-CamKIIα-eYFP or eNpHr3.0 were applied to BLA. **B**, AAV expression for animals receiving fear conditioning, extinction, and fear reacquisition with each rat represented at 10% opacity. **C**, AAV expression for animals receiving fear conditioning, extinction, and appetitive conditioning with each rat represented at 10% opacity. **D**, eYFP and eNpHR3.0 groups received extinction training with photoinhibition during shock omission. Photoinhibition augmented fear loss during extinction. A suppression ratio of 0.5 indicates no fear and a suppression ratio of 0 indicates high fear. **E**, eYFP and eNpHr3.0 groups were retrained in fear or appetitive conditioning and showed selective impairments in fear relearning. For aversive conditioning, a suppression ratio of 0.5 indicates no fear and a suppression ratio of 0 indicates high fear. For appetitive conditioning, an elevation ratio > 0.5 indicates CS elicited magazine entries. **F**, AAV-CamKIIα-eYFP or eNpHr3.0 were applied to BLA. AAV expression across all animals with each rat represented at 10% opacity. **G**, Control eYFP and eNpHr3.0 groups received extinction training with photoinhibition randomly during the inter-trial interval. Photoinhibition had no effect on extinction learning. All data are mean ± SEM and ns are listed in the main text. * p < .05.

Photoinhibition during extinction training augmented loss of conditioned fear (**Figure 4d**). Group eNpHR3.0 showed significantly less fear than group eYFP (main effect of group *F*_(1,23)_ = 12.37, *p* = .002), and fear decreased across the days of extinction (main effect of day *F*_(1,23)_ = 107.73, *p* < .001, no group × day interaction *F*_(1,23)_ = 0.91, *p* > 05). This augmentation of fear loss was temporally specific. Photoinhibition during the extinction intertrial interval (**Figure 4f**) in separate groups (eNpHR3.0, n = 5; eYFP n = 7) had no effect (main effect of trial *F*_(1,10)_ = 36.34, *p* < .001, no main effect of group and no interaction *Fs*_(1,10)_ < 1, *p* > .05) (**Figure 4g**).

Manipulations that augment extinction can produce resistance to fear relapse (Leung et al., 2012; Leung and Westbrook, 2008). To examine whether the augmented loss of fear here was accompanied by resistance to relapse, rats that had received extinction training + photoinhibition (from **Figure 4d**) were divided into four groups. Two groups (eYFP *n*=6 or eNpHR3.0 *n*=5) received fear (aversive) reacquisition training so that they again received pairings of the extinguished CS with the shock US (**Figure 4e**). There was no photoinhibition during reacquisition. The groups did not differ on the last trial of extinction training (*F*_(1,9)_ = 1.92, *p* > .05) or on the first trial of reacquisition training (*F*_(1.9)_ = 2.50, *p* > .05). However, group eNpHR3.0 was retarded in fear reacquisition (main effect of day (*F*_(1,9)_ = 18.61, *p* = .002, main effect of group (*F*_(1,9)_ = 13.99, *p* = .005, no group × day interaction (*F*_(1,9)_ < 1, *p >* .05).

Resistance to relapse might have been due to a general deficit in learning about the CS after extinction. The other two groups (eYFP *n*=8 or eNpHR3.0 *n*=6) received Pavlovian appetitive conditioning so that the extinguished fear CS was now paired with delivery of sucrose pellets. There was no photoinhibition. We measured head entries into the magazine where sucrose pellets were delivered to determine CS elevation ratios – a reliable index of Pavlovian appetitive learning (Lattal, 1999). The eNpHR3.0 group were not retarded in acquisition of appetitive learning (**Figure 4e**) (main effect of day (*F*_(1,12)_ = 19.88, *p* < .001, no main effect of group (*F*_(1,12)_ = 1.77, *p* > .05, no group × day interaction *F*_(1,12)_ = 0.55, *p* > .05). So, BLA^CaMKIIα^ optogenetic inhibition during aversive non-reinforcement enhances loss of fear during extinction and selectively impairs fear relearning.

### BLA^CaMKIIα^ photoinhibition impairs safety learning

If BLA^CaMKIIα^ optogenetic inhibition augments fear loss in an extinction paradigm, does it also augment fear reductions during safety learning? We used an AX+/BX‐ discrimination design to assess this. Rats expressing CaMKIIα-eNpHR3.0 (n = 6) or CaMKII α-eYFP (n = 7) in BLA (**Figure 5a**) received fear conditioning of the compound CS AX (i.e. CSA and CSX+) and non-reinforced presentations of the compound CS BX (CSB and CSX-) (**Figure 5b**). This establishes CSA as a fear CS. CSX is established as a weak fear CS because it is unreliably paired with the shock US. CSB is established as a conditioned inhibitor of fear (a learned safety signal) because it signals absence of the shock US (Josselyn et al., 2005; Jovanovic et al., 2012; Kazama et al., 2013; Wagner et al., 1968; Wagner and Rescorla, 1972). We photoinhibited during the expected but absent shock US on CS BX‐ trials during discrimination training only (**Figure 5b**).

**Figure 5.**
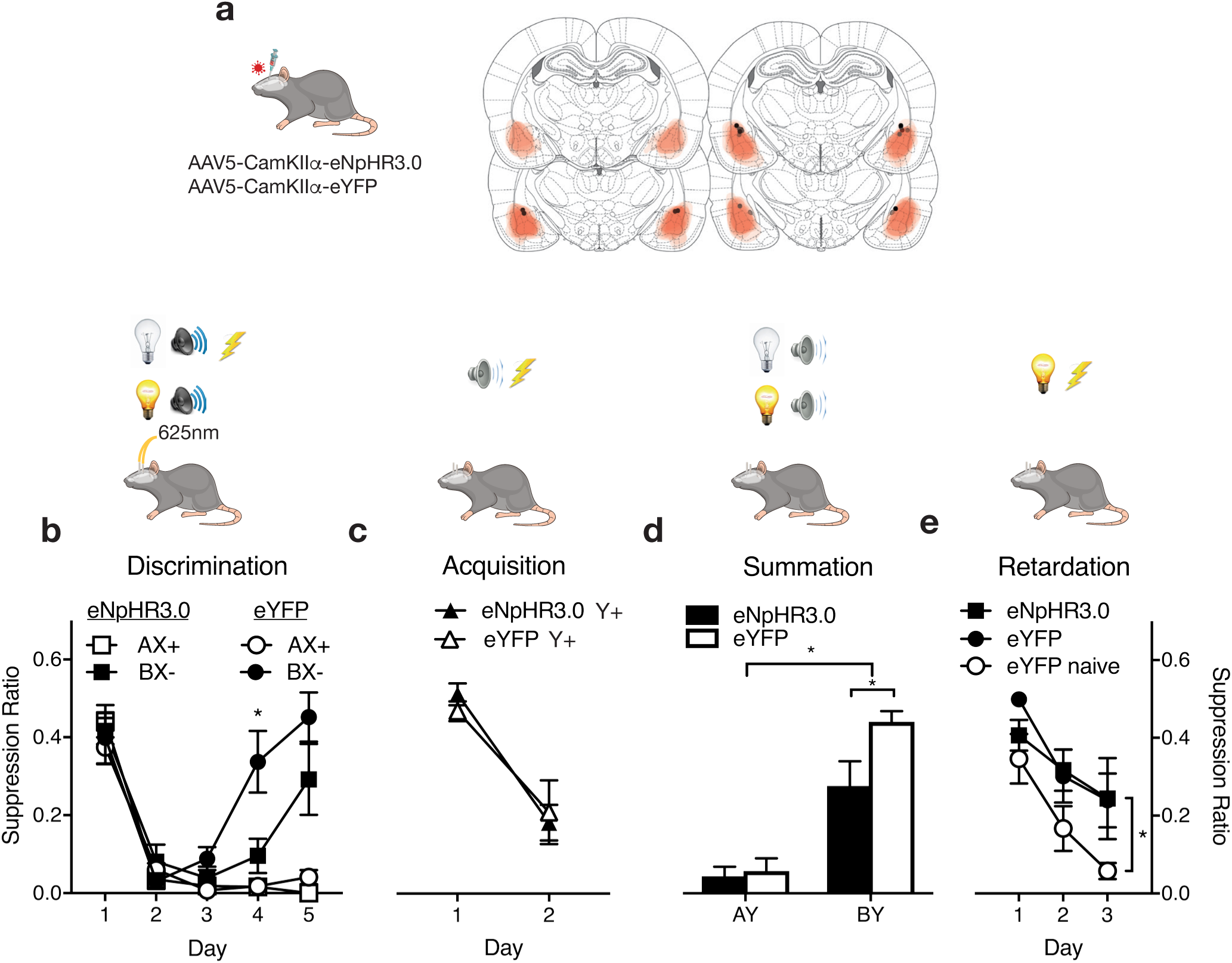
BLA^CaMKMα^ neurons and safety learning. **A**, AAV-CamKIIα-eYFP or eNpHr3.0 were applied to BLA. AAV expression with each rat represented at 10% opacity. **B**, Rats received AX+/BX‐ discrimination training to establish B as a conditioned inhibitor of fear. Photoinhibition during US omission on BX‐ trials slowed discrimination learning. **C**, Rats then received CS Y+ pairings with shock. **D**, Summation test showing poorer conditioned inhibition in the eNpHR3.0 group. **E**, Retardation test showing normal retardation in the eNpHR3.0 group. All data are mean ± SEM and *n*s are listed in the main text. * p < .05

Both groups discriminated the dangerous AX from the safe BX (**Figure 5b**). There was no main effect of group (*F*_(1,11_) < 2, p > .05), a main effect of stimulus (*F*_(1,11)_ = 29.76, p < .0001), a linear trend to AX+ (*F*_(1,11)_ = 119.20, p < .0001), and significant quadratic trend to BX‐ (F_(1,11)_ = 164.24, p < .0001). There was slower discrimination learning among eNpHR3.0 animals, day-by-day analyses of fear to BX‐ showed a significant difference on Day 4 (*F*_(1,11)_ = 6.45, p < .027).

Rats were trained to fear a new CS, Y, via pairings with shock (**Figure 5c**). CSY would serve as a target for testing the fear inhibitory properties of CSB. Both groups learned to fear CSY at the same rate (no main effect of group *F*_(1,11)_ = 0.18, p > .05; main effect of day *F*_(1,11)_ = 32.36, p < .0001, no group < day interaction *F*_(1,11)_ < 1, p > .05). This shows no long-lasting impact of photoinhibition on learning to a different CS.

The first criterion for conditioned inhibition is that CSB should reduce fear in a summation test (Rescorla, 1969) (**Figure 5d**). Rats were tested with a compound comprising two fear CSs (A and Y) versus a compound comprising the inhibitor (B) and a fear CS (Y). There was no photoinhibition. Rats should express high levels of fear to AY, because both CSA and CSY had previously been paired with shock. At issue were whether rats showed reduced fear to BY and whether this conditioned inhibition was different between the eNpHR3.0 and eYFP groups. All animals showed evidence for conditioned inhibition, less fear to BY compared to AY (main effect of stimulus: *F*_(1,11)_ = 91.74, p < .0001). However, conditioned inhibition (i.e. BY test) but not conditioned fear (AY test) was weaker in the eNpHR3.0 groups (group x stimulus interaction *F*_(1,11)_ = 5.52, p = .039; BY simple effect *F*_(1,11)_ = 6.34, p = .029; no AY simple effect *F*_(1,11)_ < 1, p > .05)

The second criterion for conditioned inhibition is that CSB should be slow to transform into a fear CS in a retardation test (Rescorla, 1969) (**Figure 5e**). We compared fear learning to CSB among the eNpHR3.0 and eYFP groups with a control group, eYFP-naive. This control group had received A+/B-discrimination training, that does not establish B as a conditioned inhibitor, and received fear conditioning to B for the first time during the retardation test. They had not received photoinhibition. All groups acquired fear during the retardation test (main effect of day: *F*_(1,16)_ = 18.88, p = .001). CSB passed the retardation test in the eNpHR3.0 and eYFP groups compared to the eYFP-naive group (*F*_(1,16)_ = 7.70, p = .014). However, unlike the summation test, the eYFP and eNpHR3.0 groups did not differ in retardation (no main effect of group or group < day interaction-*Fs*_(1,16)_ < 1, p > .05). In contrast to fear extinction, photoinhibition during aversive non-reinforcement impaired conditioned inhibition learning as measured by a summation test but, like extinction, retarded later fear relearning.

### BLA^CaMKIIα^ photoinhibition does not affect simple discrimination or latent inhibition

Simple preexposure to a CS also supports a form of inhibitory learning < latent inhibition (Lubow, 1973). Latently inhibited CSs are slow to be transformed into fear CSs but, unlike conditioned inhibitors, cannot reduce fear to other CSs (Rescorla, 1969). We asked whether BLA^CaMKIIα^ photoinhibition would affect CS learning during preexposure.

Rats expressing eNpHR3.0 (n = 5) eYFP (n = 5) in BLA (**Figure 6a**) received presentations of CSA with a shock US and separate presentations of CSB alone with photoinhibition (A+/B-). During A+/B-, discrimination training, there was a main effect for CSA across days (linear trend: F(1,8) = 48.88, p < .0001) but no main effect of group nor a group x day interaction (Fs (1,8) < 1, p > .05) (**Figure 6b**). Thus, brief photoinhibition had no effect on learning about CSB, fear learning to CSA, or simple discrimination learning. Such simple discrimination training does not normally imbue CSB with the properties of a safety signal (Rescorla and Wagner, 1972; Wagner and Rescorla, 1972) and we confirmed this by showing that CSB could not reduce fear to CSA in a summation test (no significant difference in responding to A or AB in the eNpHR3.0 and eYFP groups, Fs(1,8) < 2, p > .05) (**Figure 6c**). However, preexposure to CSB should have retarded fear conditioning due to latent inhibition. This was the case (CSA from original discrimination training v CSB, F(1,8) = 39.49, p < .0001). BLA^CaMKIIα^ photoinhibition did not strengthen or weaken this latent inhibition (no main effect of group and no interaction between factors (Fs(1,8) < 1, p > .05) (**Figure 6d**).

**Figure 6.**
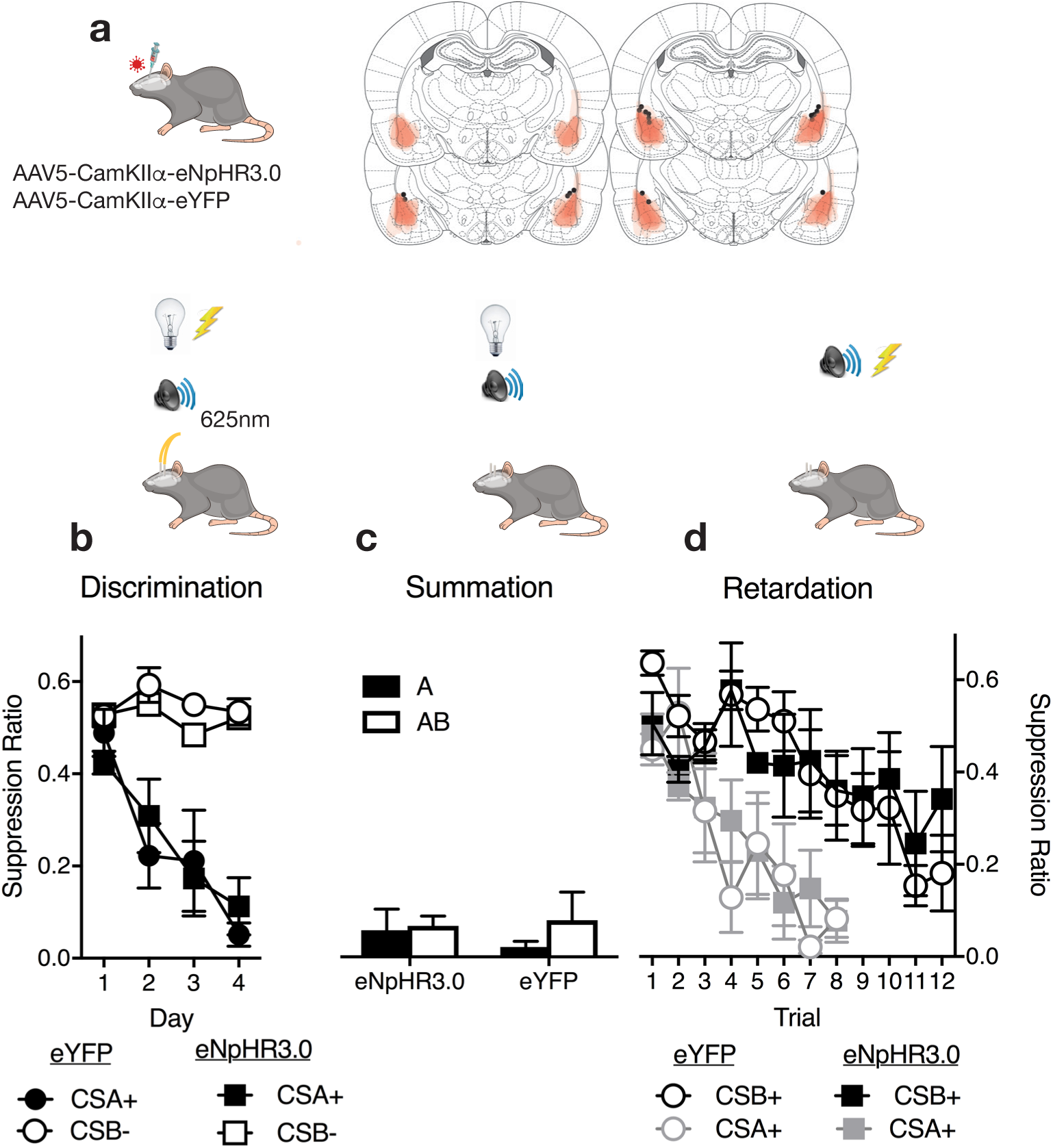
BLA^CaMKMα^ neurons and CS preexposure. **A**, AAV-CamKIIα-eYFP or eNpHr3.0 were applied to BLA. AAV expression with each rat represented at 10% opacity. **B**, Rats received A+/B-discrimination training with photoinhibition on B-trials. **C**, Summation test showing preexposure did not transform B into a conditioned inhibitor. **D**, Retardation test showing normal retardation in both eYFP and eNpHR3.0 groups. All data are mean ± SEM and ns are listed in the main text. * p < .05.

## Discussion

Here we isolated distinct features of fear, safety, and appetitive learning to understand the learning related roles of BLA^CaMKIIα^ neurons during aversive reinforcement and non-reinforcement. Photoinhibition during shock US reinforcement impaired fear learning. Photoinhibition during non-reinforcement also had effects on inhibitory learning: facilitating loss of conditioned responding during extinction, impairing relearning of fear but not learning of appetitive conditioning, and impairing learning of conditioned inhibition but not simple discrimination or latent inhibition. The most parsimonious explanation of these findings is BLA^CaMKIIα^ photoinhibition disrupts CS salience.

### Basolateral amygdala and salience

A primary function of the shock US or its absence during fear conditioning is to maintain CS salience (Hall and Rodríguez, 2017). This prevents habituation to the CS during conditioning, allowing it to be learned about and responded to. Salience is a precursor for learning, determining the eligibility of a CS to enter into an association as well as the rate at which learning occurs. It is necessary for stimulus control over behavior, determining the magnitude of conditioned responding. Highly salient stimuli have increased capacities to support learning and responding whereas low salience stimuli have these capacities reduced. The deficits in learning and responding reported here are consistent with BLA^CaMKIIα^ photoinhibition during the moments of reinforcement and non-reinforcement causing loss of CS salience.

Reductions in CS salience impair fear learning because less salient CSs are poorly learned about and less able to elicit conditioned responses. Reductions in CS salience will impair learning during extinction but also reduce conditioned responding during extinction because less salient CSs are poor at eliciting conditioned responses. In other words, the more rapid loss of fear during extinction likely reflects deficits in behavioral control by the less salient CS rather than differences in extinction learning per se. Impaired fear re-learning and responding after extinction follow from this reduction in CS salience. Impairments in safety learning occur for the same reasons. Reductions in CS salience reduce safety learning and the capacity of the safety CS to inhibit fear responses, leading to poorer performance in the summation test. Yet the same reductions in CS salience lead to typical slower fear learning and less fear responding during the retardation test.

The salience reduction had three important characteristics: it was CS-specific, long-lasting, and specific to aversive emotional states. The reduction in salience was stimulus-specific: there were no impairments of learning or responding to CSs that were not followed by BLA^CaMKIIα^ photoinhibition (e.g. offset controls, control CSs). It was long-lasting: impairments in learning and responding were detected days after last photoinhibition (e.g. retardation of fear learning following extinction or conditioned inhibition). Finally, it was specific to aversive learning, namely to CSs with a predictive relationship to the presence of shock or the absence of an expected shock. There was no impairment in appetitive conditioning to the CS or in a simple CS pre-exposure task where the CS had no such predictive relationship. This specificity to aversive learning supports distinct BLA circuits for aversive motivation (Beyeler et al., 2016; Namburi et al., 2015).

Human anxiety disorders are associated with heightened amygdala activation to threatening stimuli (Etkin and Wager, 2007). Interestingly, the consequences of BLA^CaMKIIα^ photoinhibition reported here - stimulus-specific, long-lasting, and fear-specific reductions in behavioral control - are precisely the goals of interventions for human anxiety disorders. This raises the possibility that therapeutic approaches to reduce threat stimulus salience could be especially effective in ameliorating amygdala contributions to human anxiety.

### Other BLA^CaMKIIα^ functions in fear learning

BLA^CaMKIIα^ serve a variety of other functions in fear learning, but these do not coherently account for the results reported here. Reinforcement and non-reinforcement generate a prediction error signal that instructs learning and association formation. Could the effects here be due to modulation of a Rescorla-Wagner/Temporal-Difference (Rescorla and Wagner, 1972; Sutton, 1988; Sutton and Barto, 1981) prediction error? Fear learning is instructed by positive prediction error (actual outcome of trial > expected outcome) whereas fear extinction learning is instructed by negative prediction error (actual outcome < expected outcome) (McNally et al., 2011; McNally and Westbrook, 2006). BLA^CaMKIIα^ photoinhibition could have reduced positive prediction error or augmented negative prediction error. However, conditioned inhibition depends on the same negative prediction error as extinction (Wagner and Rescorla, 1972) and should have been augmented. Instead, conditioned inhibition was impaired. Prediction error also instructs association formation indirectly by controlling attention allocated to the CS (i.e. associability) (Pearce and Hall, 1980). CS associability is upregulated when prediction error is high and downregulated when prediction error is low. BLA^CaMKIIα^ photoinhibition could have prevented upregulation of associability and so reduced fear and safety learning. However, extinction learning also involves upregulation of associability and so should have been reduced. Instead, the loss of fear during extinction was enhanced.

Activity of BLA^CaMKIIα^ during the post-training period aids consolidation of fear memories (Huff et al., 2013) and such consolidation processes can occur during the aftermath of each conditioning trial (Kamin, 1968; Wagner et al., 1973). However, extinction memories, like fear memories, undergo consolidation. If consolidation had been impaired, then fear extinction should have been impaired and later fear relearning enhanced. Instead the opposite pattern of results was observed. Finally, these effects of BLA^CaMKII^ photoinhibition cannot be due to a simple blunting of emotional reactivity. Although such a blunting could impair fear and safety learning by reducing the aversive emotional response to shock and the relief occasioned by safety, respectively, it would not lead to the long-lasting and fear specific impairments in later learning.

These roles for BLA^CaMKIIα^ in aversive prediction errors, memory consolidation, and other processes are likely embedded within distributed, multiplexed reinforcement signals (e.g., Roesch et al., 2012). They could involve distinct sub-populations of BLA^CaMKIIα^ neurons or activity of the same or different neurons at different times during the conditioning trial (Likhtik and Paz, 2015; Livneh and Paz, 2012; Sengupta et al., 2016). Regardless, our data show that the primary behavioural consequence of BLA^CaMKIIα^ optogenetic inhibition during reinforcement and non-reinforcement across fear conditioning, fear extinction, and conditioned inhibition, is a reduction in CS salience.

## Conclusion

We show that brief optogenetic inhibition of BLA^CaMKIIα^ neurons around moments of aversive reinforcement or non-reinforcement causes reductions in the salience of conditioned stimuli, rendering these stimuli less able to be learned about and less able to control fear or safety behaviours. This reduction was stimulus-specific, long lasting, and specific to aversive emotional states - precisely the goals of therapeutic interventions in human anxiety. Our findings identify a core learning process disrupted by BLA optogenetic inhibition. They suggest that a primary function of BLA^CaMKIIα^ neurons is to maintain the salience of conditioned stimuli that signal reinforcement or non-reinforcement and that this is a necessary precursor for such stimuli to gain control over fear and safety behavior.

## Acknowledgements

Supported by grants from the Australian Research Council to GPM (DP170100075 and DP160100004) and by an Australian Postgraduate Award to Y.L. Data are archived in the UNSW Long Term Data Archive (ID: D0235272).

Author contributions - Conceptualization: A.S. and G.P.M. Software: P.J.R.D.B., J.O.Y.Y., Y.L and E.Z.M. Formal Analysis: A.S., G.P.M., P.J.R.D.B., J.O.Y.Y., Y.L., E.Z.M., and J.M.P. Experimentation: A.S., P.J.R.D.B., J.O.Y.Y. and J.M.P. Writing - original draft: G.P.M and A.S. Writing - Review & Editing: All authors. Funding acquisition: G.P.M.

We thank Kathryn Baker, Kelsey Zimmerman, Nathan Holmes, and Fred Westbrook for helpful discussions of these experiments.

